# Noisy delay denoises biochemical oscillators

**DOI:** 10.1101/2023.05.17.541178

**Authors:** Yun Min Song, Sean Campbell, LieJune Shiau, Jae Kyoung Kim, William Ott

## Abstract

Genetic oscillations are generated by delayed transcriptional negative feedback loops, wherein repressor proteins inhibit their own synthesis after a temporal production delay. This delay is distributed because it arises from a sequence of noisy processes, including transcription, translation, folding, and translocation. Because the delay determines repression timing and therefore oscillation period, it has been commonly believed that delay noise weakens oscillatory dynamics. Here, we demonstrate that noisy delay can surprisingly denoise genetic oscillators. Moderate delay noise unexpectedly sharpens oscillation peaks and improves temporal peak reliability without impacting period. We show that this denoising phenomenon occurs in a variety of well-studied genetic oscillators and we use queueing theory to uncover the universal mechanisms that produce it.

## I. INTRODUCTION

Living organisms rely on periodic phenomena such as the cell cycle and circadian rhythms to survive and adapt. Biochemical interaction networks produce the periodic fluctuations in protein concentration and activity that drive such phenomena [1–3]. Negative feedback loops play an essential role in network topologies that generate periodic dynamics [4]. Generating oscillations requires a sufficient temporal delay in this feedback [4, 5], for the absence of delay results in convergence to a steady state. Delay is an inherent component of many biochemical reaction networks. It results from the sequential assembly of functional proteins via transcription, translation, folding, and phosphorylation [6–8], or arises when cytoplasmic proteins pass various obstacles and enter the nucleus to inhibit their own genes [9].

Modeling processes such as regulator protein formation and protein diffusion using explicit delays rather than intricate descriptions of the intermediate steps can be advantageous from analytical and inferential points of view [7, 10]. When introducing explicit delay representing the cumulative timing of complex processes with many stochastic intermediate steps, it is realistic to use distributed (random) delay. Nevertheless, many studies have used fixed delay for simulation and analysis [11– 14]. Oscillator studies that use fixed delay have found that fixed delay acts constructively, meaning that more delay enhances the stability of the oscillation.

By contrast, it is natural to conjecture that distributed delay weakens oscillations. This is correct in some important cases. For instance, generating strong circadian rhythms requires PERIOD proteins to enter the nucleus to inhibit their own production at a precise time of day [15–17]. The distributed delay that results from the stochasticity associated with protein generation and travel weakens the circadian rhythm [15, 18, 19]. Consequently, biological filtering mechanisms to mitigate the heterogeneous nuclear entry time have been investigated [15, 16]. Recent analysis has shown that increasing the average delay while maintaining the number of sequential processes that produce distributed delay can weaken oscillations [20].

Here, we demonstrate that distributed delay can act constructively by denoising a variety of well-studied stochastic genetic oscillators. This is surprising because distributed delay accelerates signaling in feedforward architectures [6], a phenomenon that can interfere with oscillation formation because of the need for sufficient delay in the negative feedback. We inject noise into the delay distribution by increasing the coefficient of variation (cv) while holding mean delay fixed. We find that this process sharpens oscillation peaks, thereby improving temporal peak reliability, until delay cv reaches a moderate level. Crucially, the period of oscillation remains essentially constant as this improvement occurs. We use queueing theory to uncover the universal mechanisms that drive the unexpected denoising phenomenon. The accelerated signaling induces sharper oscillation peaks, while a compensatory mechanism stabilizes the period of oscillation.

## II. RESULTS

### Mather oscillator: Denoising phenomenon

The production of mature proteins in genetic regulatory networks involves multiple sequential reactions (Fig. 1a), resulting in complex systems with numerous kinetic parameters. To simplify models of such networks, an effective delay *τ* is often used as a proxy for protein production [22–26]. Oscillatory dynamics can emerge when a mature transcription factor inhibits its own production and thereby creates a delayed negative feedback loop, provided the production delay is sufficiently large [4]. The degrade-and-fire oscillator [7] (Mather oscillator, Fig. 1b) is an important example that utilizes delayed negative feedback. It consists of a single gene that produces a repressor protein that down-regulates its own production and is cleared by dilution and enzymatic degradation. Delay is required for the production reaction, while the dilution and enzymatic degradation reactions occur instantaneously.

**FIG. 1.**
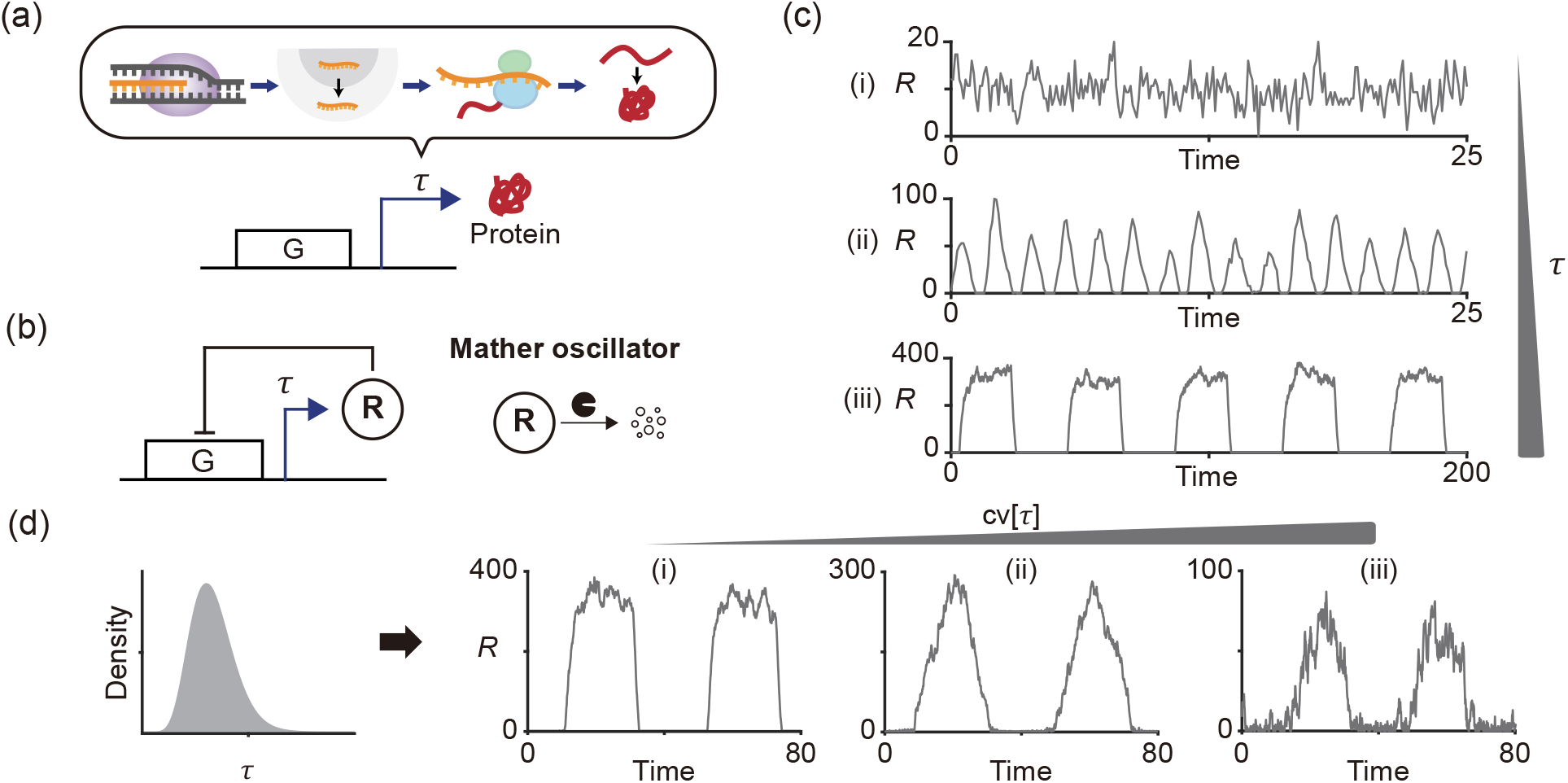
Distributed delay denoises the Mather oscillator. **(a)** Synthesizing mature regulator proteins requires a sequence of intricate processes, including transcription elongation, mRNA translocation, translation, and protein folding. Protein synthesis can be described by an effective delay, *τ*. **(b)** The Mather oscillator consists of a delayed negative feedback loop, wherein mature regulator proteins repress transcription. Regulator proteins are cleared from the system by enzymatic degradation and dilution. **(c)** When *τ* is fixed, tuning it upward induces oscillations in repressor protein level *R*(*t*). Without delay (*τ* = 0), the Mather oscillator does not oscillate (i). As the delay increases (*τ* = 0.5), oscillations emerge (ii), and a longer delay (*τ* = 20) generates strong oscillations with plateaued peaks (iii). **(d)** When *τ* follows a gamma distribution with E[*τ*] = 20, the peaks of the oscillation sharpen as cv[*τ*] increases away from zero, while the period of the oscillation appears to remain stable ((i) and (ii)). The oscillation becomes visibly noisy for large values of cv[*τ*] (iii). From (i) to (iii), cv[*τ*] values are 0.01, 0.25, and 0.49, respectively. The simulations were performed using a Gillespie-type algorithm [21].

We use a delay birth-death (dBD) framework [27] to model the Mather oscillator in particular and investigate the effect of distributed delay on stochastic oscillators in general (see Table S1 for reaction propensity functions and parameter details). We first verify that the stochastic dBD model of the Mather oscillator can generate oscillatory dynamics when the delay *τ* is fixed (i.e. cv[*τ*] = 0). The model does not oscillate when *τ* is small (Fig. 1c, (i)), while a stable oscillation emerges as the production delay increases (Fig. 1c, (ii)). As *τ* increases beyond the level at which oscillatory dynamics appear, a strong oscillation with large amplitude, long period, and plateaued peaks emerges (Fig. 1c, (iii)).

Although fixed delay has been widely used to investigate biological systems [11–14], assuming a distributed delay is more realistic due to the inherent stochasticity associated with the numerous reactions required for protein synthesis [25, 26, 28, 29]. Thus, we now investigate the impact of a distributed delay on the stochastic oscillator. We suppose *τ* follows a gamma distribution with expected value E[*τ*] = 20, the value for which fixed delay produced a strong oscillation with plateaued peaks. One might expect that delay noise decreases the period and weakens the oscillation because delay noise accelerates signaling in feed-forward genetic circuits, thereby decreasing the apparent delay [6]. Unexpectedly, however, we see that as cv[*τ*] increases away from zero, the period of oscillation does not appear to change (Fig. 1d, (i) and (ii)). Importantly, *peaks become sharper*, indicating that delay noise can denoise the Mather oscillator in terms of the temporal reliability of the oscillation peaks. Only when cv[*τ*] becomes large does delay stochasticity finally produce visible noise in repressor protein *R* timeseries (Fig. 1d, (iii)).

### Mather oscillator: Quantification of denoising

To quantify the unexpected denoising phenomenon that occurs when delay noise is moderate, we specify three quantification methods. First, we calculate the variability in the temporal distance *p* between successive peaks (green triangles in Fig. 2a, (i)) to quantify the temporal reliability of the peaks. Second, we approximate the autocorrelation function of the trajectory (*R*(*t*)),

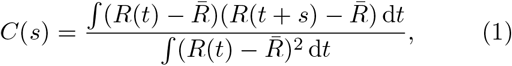

with a damped cosine function,

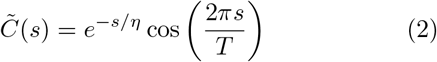

(Fig. 2a, (ii)), and then estimate the parameters *T* and *η*. Here, *T* encodes the period of oscillation, while *η* encodes how fast *C*(*s*) decays. Notice that a larger *η* implies a more robust oscillation. We use the dimensionless ratio*T/η*, referred to as oscillation noisiness, to quantify how noisy an oscillatory trajectory is. (The reciprocal *η/T* is inversely proportional to phase diffusion [30].) Finally, we quantify the peak width of each cycle (between blue triangles in Fig. 2a) by the ratio between the duration that the smoothed trajectory stays *>* 90% and *>* 10% of its maximum (Fig. 2a, (iii)).

**FIG. 2.**
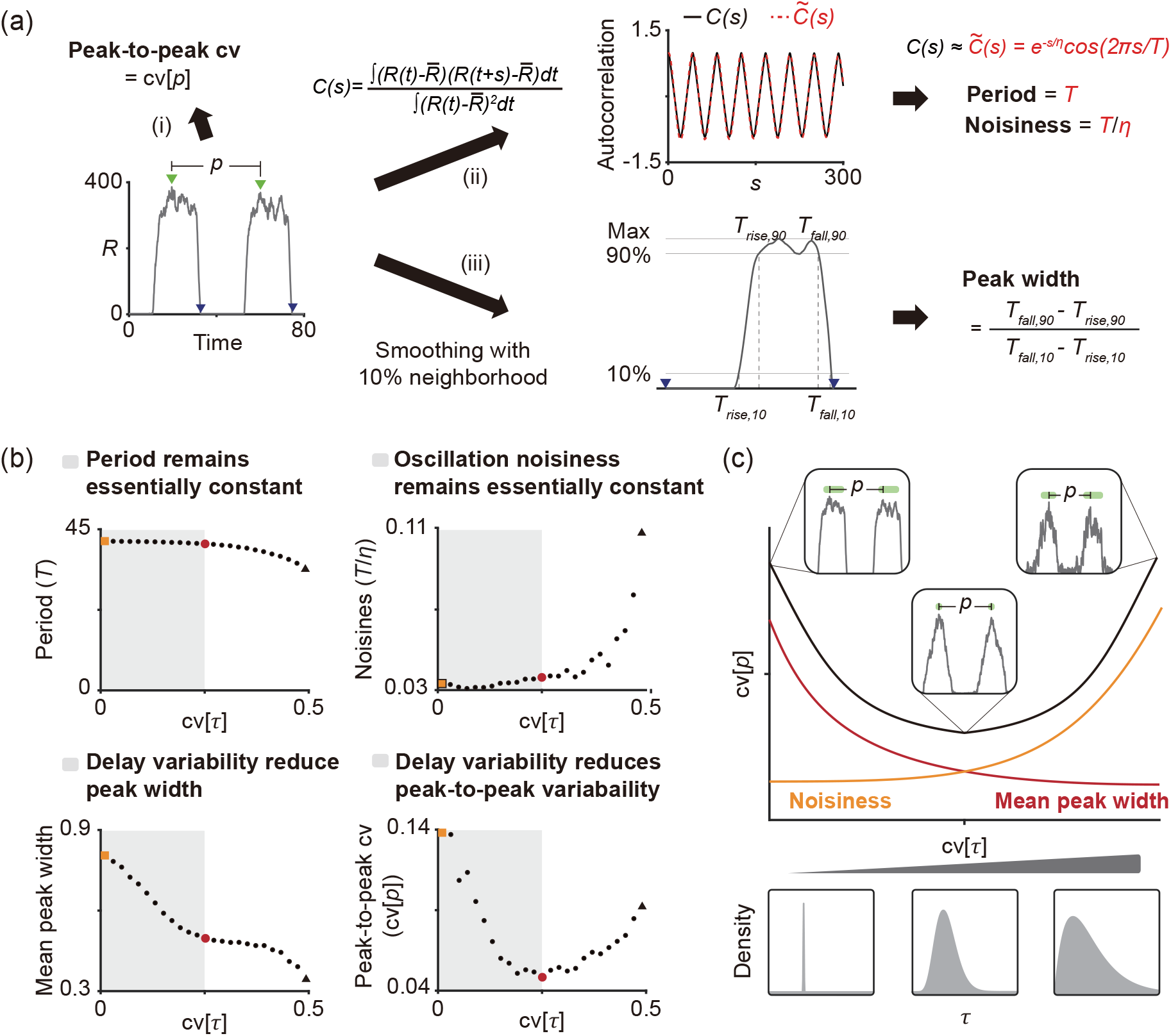
Denoising phenomenon for the Mather oscillator: Quantification and explanation. **(a)** We quantify the impact of distributed delay on the oscillation using peak-to-peak cv (cv[*p*]) (i), period (*T*) and oscillation noisiness (*T/η*) extracted from the autocorrelation function (*C*(*s*)) (ii), and peak width obtained from smoothed trajectories (iii). **(b)** Denoising occurs as cv[*τ*] increases from zero to 0.25 (gray regions), meaning that period and oscillation noisiness remain essentially constant, while peak-to-peak variability and mean peak width decrease. Peak-to-peak variability (measured using at least 500 adjacent peaks) decreases because mean peak width decreases. As cv[*τ*] increases beyond 0.25 (white regions), oscillation noisiness and peak-to-peak variability increase. The orange square, red box, and black triangle correspond to the trajectories in Fig. 1d. **(c)** The unimodal response of cv[*p*] to cv[*τ*] results from a trade-off between mean peak width and oscillation noisiness.

As cv[*τ*] increases away from zero (Fig. 2b, gray regions), period *T* and oscillation noisiness remain essentially constant. Meanwhile, mean peak width and therefore cv[*p*] decrease. When cv[*τ*] further increases (Fig. 2b, white regions), oscillation noisiness and thus cv[*p*] increase. Overall, a trade-off between mean peak width and oscillation noisiness produces the unimodal response of cv[*p*], wherein moderate delay noise yields optimal temporal peak reliability (Fig. 2c).

### Denoising biochemical oscillators: Generality and analysis

We investigate the impact of distributed delay on several additional well-studied biochemical oscillators to verify that the denoising phenomenon we have discovered is not specific to the Mather oscillator (Fig. 3). We study the Kim-Forger model [31], wherein repression is based on protein sequestration [32] and enzymatic degradation is absent; the dual-feedback oscillator [7, 33], an extension of the Mather oscillator that includes an activation loop, where the activator and repressor are cleared by coupled enzymatic degradation; and the repressilator, constructed by cyclically coupling three Mather oscillators. We model the oscillators using the dBD framework [27], as we did with the Mather oscillator. See Tables S2–S4 for model details and propensity functions. Each of the genetic oscillator models exhibits period robustness and unimodal cv[*p*] profiles across a range of values of E[*τ*] due to the trade-off between mean peak width and oscillation noisiness (Fig. 3a–d). These results suggest that distributed delay universally denoises genetic oscillators, regardless of transcriptional repression mechanism (e.g. Hill-type or protein sequestration), protein clearance mechanisms (e.g. presence of enzymatic degradation), or network structure (e.g. single or multiple feedback loops).

**FIG. 3.**
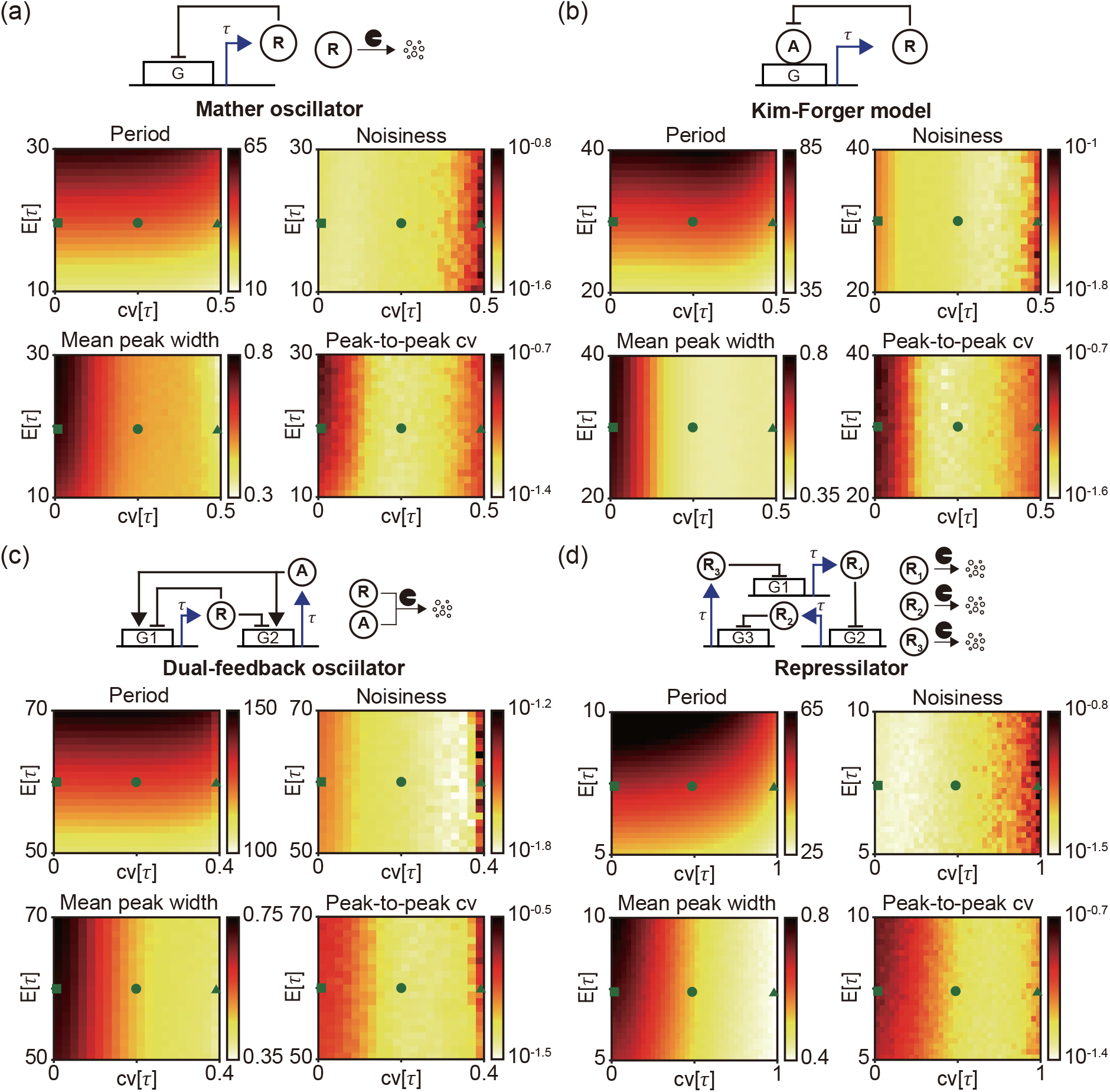
Distributed delay denoises various oscillators built upon a core negative feedback loop. We demonstrate that distributed delay denoises the Mather oscillator (a), the Kim-Forger model (b), the dual-feedback oscillator (c), and the repressilator (d). For each model, we observe denoising over a range of E[*τ*] values: As cv[*τ*] increases away from zero, period (*T*) and oscillation noisiness (*T/η*) remain essentially constant, while mean peak width and therefore peak-to-peak variability (cv[*p*]) decrease. See Fig. S1 for trajectories corresponding to green squares, circles, and triangles. The simulations were performed using a Gillespie-type algorithm [21].

We introduce and analyze a two-phase model in order to uncover the universal mechanisms that denoise genetic oscillators (see SM). The two-phase model is a simplification of the Mather oscillator that retains the core negative feedback loop. We combine stochastic analysis with deterministic techniques to show that denoising results from two effects that operate harmoniously. First, injecting noise into the delay distribution induces faster signal formation [6], reducing the number of proteins produced per cycle and thereby sharpening oscillation peaks. Second, we express the time between transcription initiation and protein clearance in a convolutional manner and show that the support of the convolution essentially does not depend on cv[*τ*]. This feature of the support explains why the period of oscillation is robust to increases in cv[*τ*]. Our analysis assumes that protein clearance via dilution can be neglected during signal formation and is therefore valid when signaling threshold is low and transcription initiation rate is high. See SM for the details that support this intuitive picture, as well as a demonstration that the analysis accurately predicts period as a function of cv[*τ*].

## III. DISCUSSION

In this Letter, we asked how distributed delay impacts the dynamics of biochemical oscillators. For a variety of well-studied genetic oscillators, we have established the counterintuitive result that injecting noise into the delay distribution sharpens oscillation peaks and thereby improves temporal peak reliability, without affecting period. We have precisely quantified this denoising phenomenon and have used queueing theory to uncover the universal mechanisms that produce it.

Sharp oscillatory peaks confer high-resolution timing upon downstream signals [34, 35], e.g. signals for plant growth [36] and starch degradation [37]. The value of sharp oscillatory peaks may extend well beyond temporal peak reliability. The mechanisms by which circadian oscillators maintain constant period over a range of temperatures remain unclear. Gibo and Kurosawa [38] argue that for circadian oscillators to compensate for temperature, it is essential that circadian waveform shapes depend on temperature. In particular, higher temperatures correspond to more nonsinusoidal waveforms. Nonsinusoidal features of neural oscillations in the brain may provide crucial physiological information related to neural communication, computation, and cognition [39]. Ongoing development of adaptive, data-driven time-frequency analysis supports the study of exotic waveforms [40].

More work is needed to fully assess the impact of distributed delay on oscillatory systems. A systematic study of network topologies would be a natural next step. Connections between network topology and oscillator robustness have been extensively examined [41, 42]. For instance, local structures that complement core topologies can significantly modulate the robustness of oscillations [41]. It would be interesting to extend such studies by including distributed delay. This could be done by introducing parameterized families of delay distributions and then studying an augmented parameter space that includes kinetic parameters and delay distribution parameters.

Beyond oscillators, the impact of distributed delay on the dynamics of biochemical systems with multiple metastable states remains to be assessed. It is known that introducing fixed delay can dramatically stabilize bistable gene networks [43]. Kyrychko and Schwartz [44] have found that broadening the width of the delay distribution reduces switching rates for a model system that admits a saddle point and a single metastable state. For stochastic systems, the interplay between distributed delay and large deviation asymptotics remains a fruitful research area.

## IV. ACKNOWLEDGMENTS

This work was supported by the Institute for Basic Science IBS-R029-C3 and Samsung Science and Technology Foundation SSTF-BA1902-01 (JKK), as well as National Science Foundation grant DMS 1816315 (WO).

## V. AUTHOR CONTRIBUTIONS

YMS and SC contributed equally to this work. JKK and WO conceptualized and designed the study. YMS conducted stochastic simulations and visualizations. SC, LS, and WO performed the mathematical analysis. YMS, LS, JKK, and WO wrote the first draft of the manuscript. All authors revised the manuscript.

## Notes

### Competing Interest Statement

The authors have declared no competing interest.

## References

[1] A. Patke, M. W. Young, and S. Axelrod, Nat. Rev. Mol. Cell Biol. 21, 67 (2020).

[2] C. Beta and K. Kruse, Annu. Rev. Condens. Matter Phys. 8, 239 (2017).

[3] A. W. Murray and M. W. Kirschner, Science 246, 614 (1989).

[4] B. Novák and J. J. Tyson, Nat. Rev. Mol. Cell Biol. 9, 981 (2008).

[5] P. Casani-Galdon and J. Garcia-Ojalvo, Curr. Opin. Cell Biol. 78, 102130 (2022).

[6] K. Josić, J. M. Lopez, W. Ott, L. Shiau, and M. R. Bennett, PLoS Comput. Biol. 7, e1002264 (2011).

[7] W. Mather, M. R. Bennett, J. Hasty, and L. S. Tsimring, Phys. Rev. Lett. 102, 10.1103/physrevlett.102.068105 (2009).

[8] A. Goldbeter, Proc. Royal Soc. B P ROY SOC B-BIOL SCI 261, 319 (1995).

[9] C. K. Macnamara and M. A. Chaplain, J. Theor. Biol. 407, 51 (2016).

[10] C. Gupta, J. M. López, R. Azencott, M. R. Bennett, K. Josić, and W. Ott, J. Chem. Phys. 140, 204108 (2014).

[11] J. Mahaffy and C. Pao, J. Math. Biol. 20, 39 (1984).

[12] L. Chen and K. Aihara, IEEE Trans. Circuits Syst. I: Fundam. Theory Appl. 49, 602 (2002).

[13] N. A. Monk, Curr. Biol. 13, 1409 (2003).

[14] K. Sriram and M. Gopinathan, J. Theor. Biol. 231, 23 (2004).

[15] S. Beesley, D. W. Kim, M. D’Alessandro, Y. Jin, K. Lee, H. Joo, Y. Young, R. J. Tomko Jr, J. Faulkner, Gamsby, et al., Proc. Natl. Acad. Sci. U.S.A. 117, 28402 (2020).

[16] S. J. Chae, D. W. Kim, S. Lee, and J. K. Kim, iScience 26 (2023).

[17] X. Cao, Y. Yang, C. P. Selby, Z. Liu, and A. Sancar, Proc. Natl. Acad. Sci. U.S.A. 118, e2021174118 (2021).

[18] J. M. Hurley, Sci. Transl. Med. 12, eabf4681 (2020).

[19] K. Montague-Cardoso, Commun. Biol. 3, 705 (2020).

[20] J. Rombouts, S. Verplaetse, and L. Gelens, bioRxiv 10.1101/2023.03.03.530971 (2023).

[21] X. Cai, J. Chem. Phys. 126, 124108 (2007).

[22] M. Barrio, A. Leier, and T. T. Marquez-Lago, J. Chem. Phys. 138, 104114 (2013).

[23] S. J. Park, S. Song, G.-S. Yang, P. M. Kim, S. Yoon, J.-H. Kim, and J. Sung, Nat. Commun. 9, 297 (2018).

[24] J. Zhang and T. Zhou, Proc. Natl. Acad. Sci. U.S.A. 116, 23542 (2019).

[25] B. Choi, Y.-Y. Cheng, S. Cinar, W. Ott, M. R. Bennett Josić, and J. K. Kim, Bioinformatics 36, 586 (2020).

[26] D. W. Kim, H. Hong, and J. K. Kim, Sci. Adv. 8, eabl4598 (2022).

[27] R. Schlicht and G. Winkler, J. Math. Biol. 57, 613 (2008).

[28] M. J. Cortez, H. Hong, B. Choi, J. K. Kim, and K. Josić, Bioinformatics 38, 187 (2022).

[29] H. Hong, M. J. Cortez, Y.-Y. Cheng, H. J. Kim, B. Choi, K. Josić, and J. K. Kim, bioRxiv 10.1101/2022.11.27.518074 (2022).

[30] Y. Cao, H. Wang, Q. Ouyang, and Y. Tu, Nat. Phys. 11, 772 (2015).

[31] J. K. Kim and D. B. Forger, Mol. Syst. Biol. 8, 10.1038/msb.2012.62 (2012).

[32] E. M. Jeong, Y. M. Song, and J. K. Kim, Interface Focus 12, 20210084 (2022).

[33] J. Stricker, S. Cookson, M. R. Bennett, W. H. Mather, S. Tsimring, and J. Hasty, Nature 456, 516 (2008).

[34] H.-H. Jo, Y. J. Kim, J. K. Kim, M. Foo, D. E. Somers, and P.-J. Kim, Commun. Biol. 1, 207 (2018).

[35] M. Foo, D. E. Somers, and P.-J. Kim, PLoS Comput. Biol. 12, e1004748 (2016).

[36] A. De Montaigu, A. Giakountis, M. Rubin, R. Tóth, F. Cremer, V. Sokolova, A. Porri, M. Reymond, C. Weinig, and G. Coupland, Proc. Natl. Acad. Sci. U.S.A. 112, 905 (2015).

[37] A. Scialdone, S. T. Mugford, D. Feike, A. Skeffington,P. Borrill, A. Graf, A. M. Smith, and M. Howard, eLife 2, e00669 (2013).

[38] S. Gibo and G. Kurosawa, Biophys. J. 116, 741 (2019).

[39] S. R. Cole and B. Voytek, Trends Cogn. Sci. 21, 137 (2017).

[40] T. Y. Hou and Z. Shi, Philos. Trans. Royal Soc. A 374, 20150194 (2016).

[41] Z. Li, S. Liu, and Q. Yang, Cell Syst. 5, 72 (2017).

[42] L. Qiao, Z.-B. Zhang, W. Zhao, P. Wei, and L. Zhang, eLife 11, 10.7554/elife.76188 (2022).

[43] C. Gupta, J. M. López, W. Ott, K. Josić, and M. R. Bennett, Phys. Rev. Lett. 111, 10.1103/phys-revlett.111.058104 (2013).

[44] Y. N. Kyrychko and I. B. Schwartz, Chaos 28, 063106 (2018).

