## Supplementary Information for "Noisy delay denoises biochemical oscillators"

### Noisy delay denoises biochemical oscillators: Supplementary Material

(Dated: May 16, 2023)

#### I. A TWO-PHASE MODEL

We present a two-phase model in order to analyze the denoising phenomenon. The two-phase model is a simplification of the Mather oscillator that retains the core negative feedback loop. We obtain it by replacing the Hill-type repression (see Table S1) with a Boolean switch and removing enzymatic degradation, yielding two propensity functions,

$$f_{\text{birth}}(R) = \begin{cases} \alpha, & \text{if } R < L; \\ 0, & \text{if } R \geqslant L, \end{cases} \tag{S1a}$$

$$f_{\text{dil}}(R) = \beta R, \tag{S1b}$$

where $L$ is a threshold that specifies the number of repressor proteins required to shut off the gene. As before, $\tau$ specifies the (random) time required to produce a mature repressor protein once transcription begins.

##### A. Phase one: Signaling time via queueing theory

We analyze the stochastic oscillation the two-phase model generates using queueing theory, a branch of stochastic analysis that has lead to numerous insights in synthetic biology [1, 2]. Phase one of the oscillation covers the time during which the production propensity is on. We assume that $L$ is low and therefore neglect dilution during phase one. We model protein production as an M/G/$\infty$ queueing system.

- **M**: Initiation of transcription (the arrival process) is memoryless. That is, the stochastic process $(I(t))_{t\geqslant 0}$ that counts the number of initiation events is a Poisson process of rate $\alpha$.
- **G**: The protein production time $\tau$ (the service process) may have a general distribution.
- **$\infty$**: In the language of queueing theory, we assume infinitely many service channels. Biochemically, this is equivalent to assuming that the availability of protein production resources is not a constraint.

Peak width is linked to the duration of phase one because we expect more initiation events when phase one is longer. Let

$$S_L = \min\{t > 0 : Z(t) = L\} \tag{S2}$$

be the signaling time at which phase one ends, where $Z(t)$ denotes the number of mature repressor proteins that the queueing system has produced by time $t$. The probability density function for $S_L$ is analytically tractable because the output process $(Z(t))$ of the M/G/$\infty$ system is an inhomogeneous Poisson process with rate $\alpha F_\tau(t)$, where $F_\tau$ is the cumulative distribution function for the delay time $\tau$. It follows that the probability density function for $S_L$ is given by

$$f_{S_L}(t) = \frac{\bar{Z}(t)^{L-1}}{(L-1)!} e^{-\bar{Z}(t)} \frac{\mathrm{d}\bar{Z}(t)}{\mathrm{d}t}, \tag{S3}$$

where

$$\bar{Z}(t) \equiv \mathrm{E}[Z(t)] = \alpha \int_0^t F_\tau(s)\,\mathrm{d}s. \tag{S4}$$

The mean and variance of $S_L$ are therefore

$$\mathrm{E}[S_L] = \int_0^\infty \frac{s^{L-1}e^{-s}}{(L-1)!}\bar{Z}^{-1}(s)\,\mathrm{d}s, \tag{S5}$$

$$\mathrm{Var}[S_L] = \int_0^\infty \frac{s^{L-1}e^{-s}}{(L-1)!}(\bar{Z}^{-1}(s))^2\,\mathrm{d}s - (\mathrm{E}[S_L])^2. \tag{S6}$$

Importantly, the expected signaling time $\mathrm{E}[S_L]$ is a *decreasing* function of $\mathrm{cv}[\tau]$ [1]. Intuition-building asymptotic expressions have been computed for the case of Bernoulli delay. Suppose that $\tau$ takes two values, $\mu_\tau - \sigma_\tau$ and $\mu_\tau + \sigma_\tau$, with equal probability. In this Bernoulli case, we have

$$\mathrm{E}[S_L] \approx \mu_\tau + \frac{L}{\alpha} \tag{S7}$$

when $\sigma_\tau$ is small, and

$$\mathrm{E}[S_L] \approx \mu_\tau + \frac{L}{\alpha} - \big(\sigma_\tau - \frac{L}{\alpha}\big) \tag{S8}$$

when $\sigma_\tau$ is large. The transition point between these two regimes is $\sigma_\tau \approx L/\alpha$. Intuitively, $\mathrm{E}[S_L]$ becomes sensitive to $\sigma_\tau$ when the order in which events exit the queue typically no longer matches the order in which they entered the queue. See Fig. S2 for an illustration of this skipping effect.

We expect $\mathrm{E}[S_L]$ to depend sensitively on $\sigma_\tau$ for any delay distribution when

$$\sigma_\tau \geqslant \frac{L}{\alpha}. \tag{S9}$$

This is precisely the parameter regime that motivates our analysis of the two-phase model ($L$ low, $\alpha$ large). See Fig. 2 of [1] for an illustration of the sensitivity of $\mathrm{E}[S_L]$ to $\sigma_\tau$ when $L$ is low and $\tau$ is gamma-distributed.

Since $\mathrm{E}[S_L]$ decreases as $\mathrm{cv}[\tau]$ increases, it follows that for the two-phase model, mean peak width decreases as $\mathrm{cv}[\tau]$ increases away from zero. This peak sharpening occurs because the expected number of proteins queued for production, $\mathrm{E}[I(S_L)] = \alpha\,\mathrm{E}[S_L]$, also decreases as $\mathrm{cv}[\tau]$ increases, reducing the expected amount of time that $R$ will fluctuate around the plateau level $\alpha/\beta$. (The plateau level $\alpha/\beta$ is the equilibrium level at which the maximal queue outflow rate $\alpha$ and dilution balance.)

The fact that delay noise induces quicker signaling explains the peak-sharpening phenomenon we have observed in the Mather oscillator, the Kim-Forger model, the dual-feedback oscillator, and the repressilator.

##### B. Phase two: Queue outflow

Since delay noise induces quicker signaling, it is surprising that for the four oscillators we have tested, period $T$ remains essentially constant as $\mathrm{cv}[\tau]$ increases away from zero. We explain this robustness by analyzing the second phase of the two-phase model. Phase two begins at time $S_L$ and ends after the ensuing peak, when $R$ returns to level $L$.

We analyze phase two using a deterministic queue outflow approach. Such an approach is sensible because $\alpha$ is assumed to be large. Let $Q(t)$ denote the number of molecules in the queue at time $t$. We develop a first-order model for queue outflow by conditioning on $S_L$ and $Q(S_L)$. Suppose that $S_L = s_L$ and $Q(S_L) = N$. These assumptions imply that $I(s_L) = N + L$. Since the initiation process $I$ is a Poisson process, it follows that these $N + L$ initiation events are distributed as the order statistics of $N + L$ independent random variables, each uniformly distributed on $[0, s_L]$.

Let $U$ be a random variable with the uniform distribution on $[0, s_L]$ and let $f_U$ denote the probability density function for $U$. The convolution $f_U * f_\tau$, when multiplied by $N + L$, gives the queue outflow rate in our queue outflow model,

$$\dot{X} = (N+L)(f_U * f_\tau)(t) - \beta X \tag{S10a}$$

$$X(s_L) = L, \tag{S10b}$$

where $X(t)$ denotes the number of mature proteins that have exited the queue but have yet to be cleared by dilution.

Let $t_a$ be the time at which the count of mature proteins returns to $L$, obtained by solving (S10):

$$L = \exp(\beta(s_L - t_a))\left(\int_{s_L}^{t_a}(N+L)(f_U * f_\tau)(w)e^{\beta(w-s_L)}\,\mathrm{d}w + L\right). \tag{S11}$$

We obtain an analytical prediction $T_a$ for the period of the two-phase model by integrating away the conditioning on $S_L$ and $Q(S_L)$:

$$T_a = \int_0^\infty \Big( \sum_{N=0}^\infty t_a(s_L, N) p_{s_L}(N) \Big) f_{S_L}(s_L) \mathrm{d}s_L. \tag{S12}$$

Here $p_{s_L}$ is the probability mass function for the number of proteins that remain in the queue at the signaling time $S_L$, conditioned on knowing that $S_L = s_L$. The analytical framework accurately predicts the period of the two-phase model (Fig. S2).

The mean period of the two-phase model remains stable as $\mathrm{cv}[\tau]$ increases away from zero (Fig. S2f). This is consistent with the first four oscillators we have tested, suggesting that a core delayed negative feedback loop induces stability of period. The nature of the convolution $f_U * f_\tau$ intuitively explains why delayed negative feedback induces such stability. Let $\hat{U}$ be a random variable with the uniform distribution on $[0, \mathrm{E}[S_L]]$ and let $f_{\hat{U}}$ be the corresponding pdf. When $\mathrm{cv}[\tau]$ is small, the right tail of $f_\tau$ (which governs the length of the queue outflow phase) is narrow, while the support of $f_{\hat{U}}$ (which governs the length of the signaling phase) is broad (Fig. S2c). As $\mathrm{cv}[\tau]$ increases, the right tail of $f_\tau$ moves rightward, but the support of $f_{\hat{U}}$ compensates by moving leftward (Fig. S2d). This compensatory mechanism is balanced in the sense that the time at which the cumulative distribution function of $f_{\hat{U}} * f_\tau$ essentially reaches one (a proxy for the period of the two-phase model) is not sensitive to $\mathrm{cv}[\tau]$ (Fig. S2e). Consequently, the period of the two-phase model remains stable (Fig. S2f).

#### C. Extensions

Our analysis of the two-phase model can be extended in many ways. More sophisticated queueing systems can be used to model phase one when resource limitations during protein production impact production dynamics. Stochastic processes outside of the Poissonian framework describe bursty protein production. Phase two can be modeled stochastically when molecule counts are low to moderate.

#### II. SUPPLEMENTARY FIGURES

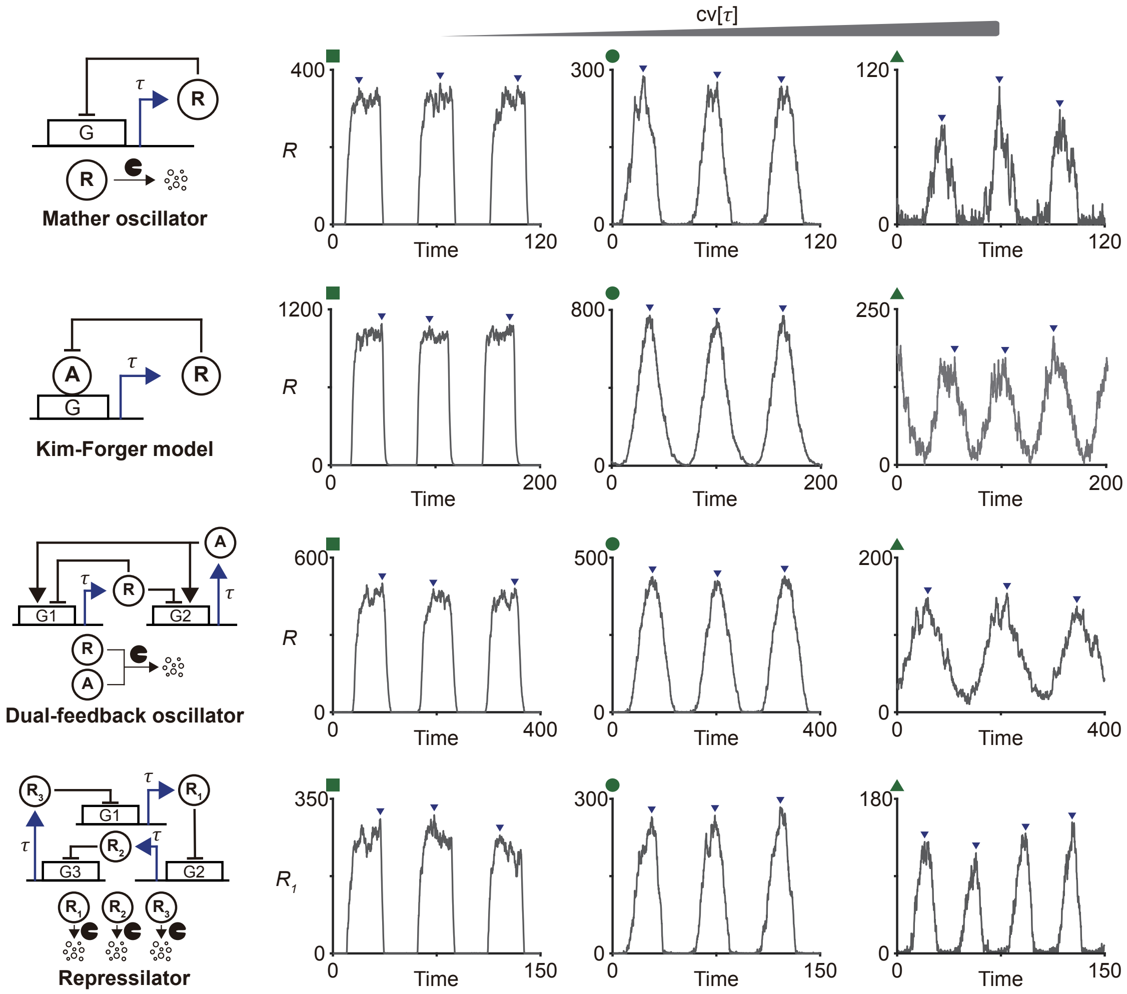


FIG. S1. **Distributed delay denoises various oscillators built upon a core negative feedback loop.** For each system, we plot a representative oscillatory trajectory for each of three values of cv[$\tau$] (see Fig. 3). Peak-to-peak distance is given by temporal distance between consecutive blue triangles. When cv[$\tau$] is small (green squares), the oscillatory trajectories exhibit plateaued peaks, resulting in highly variable peak-to-peak distances. Oscillation peaks sharpen as cv[$\tau$] increases (green circles), thereby reducing the variability of peak-to-peak distances. The oscillatory trajectories become visibly noisy when cv[$\tau$] grows sufficiently large (green triangles). See Tables S1-S4 for the corresponding propensities and parameters.

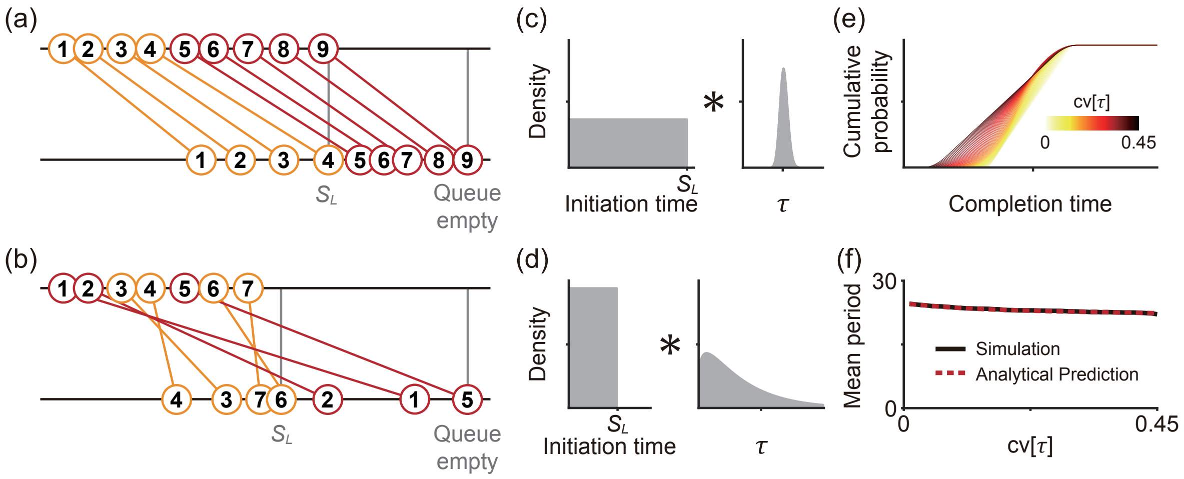


FIG. S2. **For the two-phase model, temporal compensation between phases one and two explains why the period remains stable as delay variability increases. (a,b)** Cartoons of queue dynamics when cv[$\tau$] is (a) low and (b) moderate. Here the signaling threshold is $L = 4$. When cv[$\tau$] increases from low to moderate, signaling becomes faster (E[$S_4$] decreases), but the queue requires more time to empty after the signaling threshold is crossed. These effects balance, resulting in a stable period. **(c,d)** The convolution $f_U * f_\tau$ in (S10) governs the analytical prediction for period. We plot the pdf of $\tau$ and representative pdfs of $U$ for cv[$\tau$] low (c) and moderate (d). As cv[$\tau$] increases, the support of the representative pdf of $U$ narrows, while the tail of the pdf of $\tau$ grows heavier. **(e)** This compensatory mechanism is balanced in the sense that the time at which the cumulative distribution function of $f_{\hat{U}} * f_\tau$ essentially reaches one (a proxy for the period of the two-phase model) is not sensitive to cv[$\tau$]. **(f)** An analytical prediction for the mean period of the two-phase model closely matches the results of simulations. System configuration for (e) and (f): $\alpha = 1000$, $\beta = 1$, and $L = 10$; $\tau$ is distributed uniformly with E[$\tau$] = 10 and cv[$\tau$] between 0 and 0.45.

#### III. SUPPLEMENTARY TABLES

TABLE S1. Propensities and parameter values for the Mather oscillator [3]. Bold font denotes the reaction with delay.

| Reactions | Propensities | Parameters |
|---|---|---|
| $\boldsymbol{\emptyset \to R}$ | $\alpha \frac{C_0^2}{(C_0+R)^2}$ | $\alpha = 300,\ C_0 = 10$ |
| $R \to \emptyset$ | $\gamma \frac{R}{R_0+R}$ | $\gamma = 106,\ R_0 = 0.0001$ |
| $R \to \emptyset$ | $\beta R$ | $\beta = 0.6$ |

TABLE S2. Propensities and parameter values for the Kim-Forger model [4]. The mRNA transcription and protein translocations of the original model have been replaced with a distributed delay. Bold font denotes the reaction with delay.

| Reactions | Propensities | Parameters |
|---|---|---|
| $\boldsymbol{\emptyset \to R}$ | $\frac{\alpha}{2}(X + \sqrt{X^2 - 4A_T K_d})$, $X = A_T - R - K_d$ | $\alpha = 10,\ A_T = 100,$ $K_d = 0.001$ |
| $R \to \emptyset$ | $\beta R$ | $\beta = 1$ |

TABLE S3. Propensities and parameter values for the dual-feedback oscillator [3, 5]. Bold font denotes reactions with delay.

| Reactions | Propensities | Parameters |
|---|---|---|
| $\mathbf{\emptyset \to R}$ | $\alpha \frac{f^{-1}+c^{-1}A}{(1+c^{-1}A)(1+c^{-1}R)^2}$ | $\alpha = 300,\ f = 2,\ c = 50$ |
| $R \to \emptyset$ | $\gamma \frac{R}{r+R+A}$ | $\gamma = 40,\ r = 1$ |
| $R \to \emptyset$ | $\beta R$ | $\beta = 0.2$ |
| $\mathbf{\emptyset \to A}$ | $k\alpha \frac{f^{-1}+c^{-1}A}{(1+c^{-1}A)(1+c^{-1}R)^2}$ | $k = 2,\ \alpha = 300,$ $f = 2,\ c = 50$ |
| $A \to \emptyset$ | $\gamma \frac{A}{r+R+A}$ | $\gamma = 40,\ r = 1$ |
| $A \to \emptyset$ | $\beta A$ | $\beta = 0.2$ |

TABLE S4. Propensities and parameter values for the repressilator. Bold font denotes the reactions with delay.

| Reactions | Propensities | Parameters |
|---|---|---|
| $\mathbf{\emptyset \to R_i}$ | $\alpha \frac{C_0^2}{(C_0+R_{i+1})^2}$, $(i = 1, 2, 3; R_4 = R_1)$ | $\alpha = 180,\ C_0 = 10$ |
| $R_i \to \emptyset$ | $\gamma \frac{R_i}{R_0+R_i},\ (i = 1, 2, 3)$ | $\gamma = 80,\ R_0 = 1$ |
| $R_i \to \emptyset$ | $\beta R_i,\ (i = 1, 2, 3)$ | $\beta = 0.4$ |


[1] K. Josić, J. M. López, W. Ott, L. Shiau, and M. R. Bennett, PLoS Comput. Biol. **7**, e1002264 (2011).
[2] W. H. Mather, J. Hasty, L. S. Tsimring, and R. J. Williams, Biophys. J. **104**, 2564 (2013).
[3] W. Mather, M. R. Bennett, J. Hasty, and L. S. Tsimring, Phys. Rev. Lett. **102**, 10.1103/physrevlett.102.068105 (2009).
[4] J. K. Kim and D. B. Forger, Mol. Syst. Biol. **8**, 10.1038/msb.2012.62 (2012).
[5] J. Stricker, S. Cookson, M. R. Bennett, W. H. Mather, L. S. Tsimring, and J. Hasty, Nature **456**, 516 (2008).
